# Dead or Just Asleep? Variance of Microsatellite Allele Distributions in the Human Y-Chromosome

**DOI:** 10.1101/008227

**Authors:** Joe Flood

## Abstract

Several different methods confirm that a number of micro-satellites on the human Y-chromosome have allele distributions with different variances in different haplogroups, after adjusting for coalescent times. This can be demonstrated through both heteroscedasticity tests and by poor correlation of the variance vectors in different subclades. The most convincing demonstration however is the complete inactivity of some markers in certain subclades – “microsatellite death”, while they are still active in companion subclades.

Many microsatellites have declined in activity as they proceed down through descendant subclades. This appears to confirm the theory of microsatellite life cycles, in which point mutations cause a steady decay in activity. However, the changes are too fast to be caused by point mutations alone, and slippage events may be implicated.

The rich microsatellite terrain exposed in our large single-haplotype samples provides new opportunities for genotyping and analysis.

## Introduction

Microsatellites or Short Tandem Repeat (STR) sequences are very widely used genomic tools. The counts of numbers of repeats of the base motif sequence in each of a set of microsatellite markers provide phenotyping information with a good degree of discriminatory power, because microsatellites mutate orders of magnitude more frequently than other parts of the chromosome. For this reason they are extensively applied in evolutionary studies, medical genetics, anthropological, genealogical, population and forensic research [1]. Bhargava and Fuentes [2] state, “the ease of use, high reproducibility, low cost and abundance of microsatellites in living organisms makes them ideal markers for genetic analysis”.

Despite the importance of microsatellites to different disciplines, much of the literature on changes in their counts and their differences within different populations has frequently involved small samples and small numbers of markers, which has possibly concealed more complex behaviours of the microsatellites.

In the standard stepwise mutation model which views microsatellite change as a symmetric random walk, it is presumed that microsatellite markers always mutate independently at the same rate, increasing or decreasing the current count randomly in a single step of constant probability. While a number of authors have tried to relax these conditions to establish a more generalised model (see Bird [3]), by and large the fundamental assumption that each marker has a unique constant mutation rate which has no temporal or population variation has been largely unchallenged. Bird writes “There is no reason to believe that mutational behaviour for STRs will differ between male Y-haplogroups or geographic regions”.

We show that this is not the case; that Y-microsatellites commonly differ in their mutational behaviour between haplogroups and over time. In particular we show that some STRs eventually become inactive, no longer mutating at all, invalidating Sun *et al.*’s [4] claim that microsatellites “are essentially guaranteed to be polymorphic.”

## Stationary markers

The observation that mutational probabilities can slow with time emerged from an investigation into why different Y-haplogroups can be distinguished by the value of particular microsatellite markers, which is not so easily explained in the standard stepwise model. Haplogroup formation depends on point mutations and is independent of microsatellite values, so the distribution of microsatellite allele count values in descendants through any fixed stochastic process might also be expected to be identical. However the modal marker values can be very different, allowing unique “microsatellite signatures” for many haplogroups. It was discovered that this discrimination is due in part to marker senescence and eventual “microsatellite death” in which no mutational activity occurs (unpublished data).

Consider for example the subclade R-L48+, to which about 20 per cent of the male population in the vicinity of the North Sea belongs. Our “convenient sample” [5], which is taken from online projects at a commercial testing company, has 286 individuals with distinct haplotypes who have tested 111 microsatellite markers, and a further 330 who have tested 67 markers. All have tested positive for the L48 point mutation which defines the subclade.

One of the standard markers, DYS593, takes a constant allele value of #15 in the sample. In smaller samples this could be due to chance, but not in samples of this size. R-L48 has a coalescence time of at least 98 generations, as we show later in the paper. Burgarella and Navascuez [6] estimate the mutation rate for DYS593 to be *p* = 4.387 × 10^−4^ per generation, so the probability that the marker will not mutate at all in a line from the founder to a single modern individual over 98 generations is at most *(1-p)^98^* or 0.958.

For medium sized samples, it is quite likely that some slow marker will not show polymorphism. With a sample size of 105, we could expect there to be no variance of the marker above in about one per cent of samples. So for a hundred or so similar markers with this same low mutation rate, one of them would likely show as being invariant in such a sample.

However the probability of invariance over the whole sample for an active marker rapidly declines with sample size, and with any sample size over about 150, the probability of invariance becomes increasingly small. With no variance in a sample of 286 men, the chance is about one in 5 million that the marker is active.

This argument might possibly be invalidated if the R-L48 sample contained a dominant group with a much more recent coalescent time, but we find no evidence of this. Alternatively, if the mutation rate is more than an order of magnitude lower than has been stated here a zero sample variance could result: *p* = 2.4 × 10^−5^ would given an even chance of a sample of 286 men having no variance in some marker. But in fact in an alternative estimate, Ballantyne *et al.* [7] find the mutation rate for DYS593 is about five times as great as the rate we have used (see table A1), so the analysis above is based on the lowest available estimate.

But when we consider the remainder of the parent subclade of R-L48, R-U106 + L48-, for which we have a sample of similar size, we find that the variance in DYS593 is low but more than the variance in about four other slow moving markers (see table A1 in the accompanying material). It is virtually certain that this marker is inactive within R-L48, but not within other subclades of R1b.

The second constant marker DYS472 within R-L48+, has been inactive over a much longer period of time; and almost in almost every haplogroup it takes the value #8. The only known variations in the marker are in a vestigial ancient A1a cluster found in Northern Europe, where it is fixed at #9, and as a variant within the E1b1b1 haplogroup, where it takes values #9 and #10. A few other isolated cases are known of mutation in DYS472, but these could well be testing errors. The marker is to all intents and purposes fixed, and when mutation occurs, it is effectively a unique-event polymorphism.

Jobling (personal communication) has suggested that the low allele count of this marker may be responsible for its apparent inactivity. However this is still within the range for polymorphism – other markers such as DYS450, DYS459, and DYS590 take this or even lower values and still show limited polymorphism.

Other markers in R-L48 also show a much lower variance than might be expected. DYS426 and DYS436 have only four men in samples of over 550 who do not have the modal allele – suggesting that these markers are probably inactive in a substantial part of the subclade. These are slow markers, but the faster marker DYS556 has only 3 deviations among 255 men – virtually impossible by chance.

Where there is one invariant marker, there may be many. We have not comprehensively surveyed all the haplotypes to search for microsatellite marker invariance; however we have found subclades of haplogroup R1a, haplogroup E and haplogroup C with markers that appear to be inert: see Table 1.

**Table 1.** Suspected inert markers, selected subclades

| *Subclade* | *Mean*<br>*subclade*<br>*variance\** | *Inert markers (sample size in brackets)* |
| --- | --- | --- |
| R-L48 | 0.213 | DYS593 (286) |
| R1a1a1 | 0.278 | DYS436 (1089) DYS636 (459) |
| E1b1b1c | 0.615 | DYS590 (354), DYS492 (354),<br>DYS636, DYS425 both null (354), |
| C3 | 0.650 | DYS426 (455) DYS472 (138) |

Note (\*) The mean variance should be proportional to the number of generations to coalescence, by the standard model. DYS472 is invariant everywhere and has been excluded.

**Table 2.**
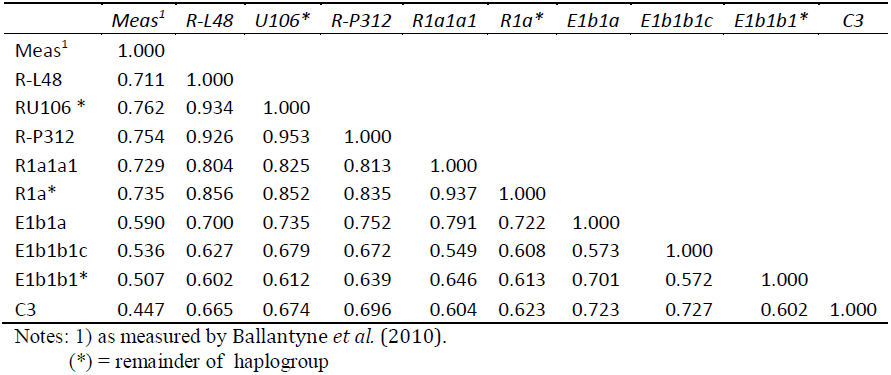
Correlations between 111 marker variances and with measured mutation rates in selected subclades.

The marker DYS636 within haplotype R1a supports the proposition that marker activity may deteriorate as an aging microsatellite proceeds down through subclades. This marker is moderately active in other haplogroups, but in the earlier subclades of R1a it shows very limited activity with only 5 deviations in 257 records. In the haplotypes descended from R1a1a (R-M417) it is inactive.

Haplotype E1b1b1c (E-M123), which is found in a patchy fashion throughout Western Asia, Ethiopia and eastern Europe, is a stand-alone subclade with peculiar characteristics that suggest it may have been formed through some a significant Y-chromosome alteration. One marker (DYS492) gives a null result for all members of the subclade, suggesting that this microsatellite has been badly damaged by a full or partial deletion. At least another four markers are inert in E-M123. It has a large genetic distance from other parts of the old E1b1b1 E-M35 clade (formerly known as E3b) to which it belongs, with different modal values on 37 out of 111 markers.

Another interesting case is the C haplogroup, which is the earliest in Eurasia (our sample consists almost entirely of the branch C3). Because of the great age of the haplogroup, many Y-STRs have larger variances in C3 than in more recent haplogroups. However, two markers DYS426 and DYS472 are inert in C3, and possibly several more.

### Different mutation rates by subclade and increasing senescence

Although it has always been considered that the statistical distribution of individual microsatellite markers within subclades is identical and affected only by the number of generations since the common ancestor, an inspection of the actual distributions shows something very different. Little (2007) placed a table of allele frequencies for 67 markers for 1866 men in eight Y-haplogroups on a public website. The distributions for markers differ between haplogroups not just in modal values, but in general structure – skewed to a greater or lesser extent and sometimes in different directions, with a greater or lesser spread to the mode. Our own larger samples confirm this. Little also presents a table of variances which differ a great deal between clades on many markers. We show the same in Table A1, on samples from the same source as Little but with seven years of further DNA contributions from the public.

Now there are several reasons why the distributions might be different for a given locus. The results might simply be the results of sample variance (stochasticity). Or they may be actual differences in the distribution - caused by founder effects or through more profound unique event polymorphisms such as medium or large inserts or deletions. Both these can cause changes in means and variances of microsatellites. Pointwise mutations will not change mean values – but as we shall see they can change the rate of mutation and hence the variances of the distributions.

With the large samples of up to 3000 that we now have for individual haplogroups, it becomes clear that the observed differences in variances (see Table A1) are not the result of sampling error. With samples of only 100, the F test shows that a ratio of 1.4 between the variances of two samples is significant at the 95% confidence level, and a ratio of 1.6 is sufficient at the 99% confidence level. With 300 in the samples the ratios for 99% significance are 1.21 and 1.31 respectively. Many markers exceed these tolerances in different haplogroups.

We also ran tests for homogeneity of variance (Levene’s test) which suggested on about 15 markers within R-L48+ that microsatellite distributions were heteroscedastic with respect to the neighbouring R-L48-groups within the parent R-U112 (Northern European R1b). As well as this, about thirteen more markers have inherited low variances from predecessor subclades of R1. We can therefore observe deteriorated markers (ones in which the pure repeat structure is damaged) being passed to subclades, so that correlation between variations in samples with a common heritage exists.

A moire elegant way of showing this inheritance is to look at the statistical correlations of variance vectors. If the single probability model were sufficient then the variances ought to be closely correlated between samples from different subclades. The variances in each sample should be 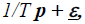 where *T* is the coalescent time for that sample, ***p*** is the vector of mutation probabilities per generation, and 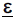 is a random error vector. The correlation matrix of variances for the different subclade samples in Table A1 is shown in table A2.

If the single probability model holds, with ***p*** constant for all samples, and the differences in variances between subclade samples are purely random, then the correlations of variances in separate haplogroups should be close to 1, should also correlate closely with measured mutation rates, and otherwise should have no discernable pattern. Instead the table shows a great deal of structure:

- We have used measured transition probabilities from [7] in Table A1 rather than from [6] because the correlations with most haplogroups are about 15 per cent higher than if probabilities from the latter were used. There are alternative values for about 30 markers which could improve the correlations by a further 15 per cent.
- The R-L48 sample correlates most closely with its nearest genetic neighbours: the rest of R1b, then R1a, then E and C.
- The correlation between variances for markers in E1b1a with other parts of E1b falls as the genetic distance increases.
- Subclades of E correlate quite poorly to each other – less than they do with C3. This is partly due to bimodal distributions, which cause variances for some markers to be large. As these haplotypes do not subsequently converge or overlap as is the case for recent R1b subclades, the bimodality may well be created by ancient unique events rather than by bottlenecking and founder effects.

The evidence therefore points firmly away from the constant probability random walk model. If we accept the different variances by haplotype in Table A1 are real, then we are looking at founder effects or at changes in mutation probabilities in the different haplogroups.

Founder effects are in fact evident. Many of the larger variances in Table A1 – those over 1.8 or so which tend to dominate the average – are the result of skewed or bimodal distributions. A closer examination of most of the skewed distributions shows that they are actually the union of two or more similar normal distributions with different modes and coalescent times; generally caused by a founder effect which changes the point at which the random walk starts for a significant part of the sample (see [8] for a clear exposition). If the two means are very different so that the combined distribution is clearly bimodal, there is a suspicion that larger structural changes to the microsatellite have taken place, not just bottlenecking. In any case, the measured variance of the distribution becomes much larger in the presence of bimodality – and even a single bimodal marker can impact heavily on coalescent time estimation.

Bimodality is not an issue in the markers showing very low variances. Lack of spread (in particular, the very high kurtosis shown by many markers) can be taken as a sign of deteriorating microsatellite function, in the sense of the “life-cycle” microsatellite model we discuss later. Within haplogroup R1b, as we have seen, there is evidence of a chain of decreasing marker activity in more recent subclades. In Table A1, 23 markers are showing lower variance than expected within R-L48. Now ten of these low-variance markers are peculiar to R-L48, four occur throughout the parent R-U102, three more fall across the whole “Western R1b” R-M239 haplotype, and a further six occur in R1a haplotype as well, suggesting a clear process of senescence from the R1-founder through to descendant haplotypes.

A similar decline occurs in E1b haplogroup. In E1b1b1c, 35 markers have low variances, of which 12 are low throughout the E1b1 clade, a further 9 are low in the intermediate subclade E1b1b1, and 14 are particular to E1b1b1c.

### Dating the haplogroups

An approximate coalescence time (TMRCA) is needed in order to confirm a marker is inactive. There are several means of estimating TMRCA for a group of men, of which the most favoured has been to use the average sample variance. The variance quotient method is extremely simple – one divides the average (or total) variance on a set of markers by the average (or total) of the marker mutation rates. There has been a very large literature since Goldstein *et al.* [9] introduced this method, and it is claimed to be a very reliable unbiased estimator of coalescent times by its adherents ([3], [4]).

Because it takes the square of deviation from the mean, the variance is not a robust measure as it is sensitive to outliers and to measurement and processing errors. The method is unreliable for other reasons, especially the very large spread on the 95% confidence limits for mutation rates ([6], Table 1, place a hundred-fold confidence range on their probability estimates).

However the source of error that is our main concern is that mutation slippage rates estimated from general populations have been used within particular haplogroups which have different mutation rates. For example, using different sets of markers in a sample from a specific haplogroup can give very different estimates of coalescent times [10] – because a number of the markers have very different mutation rates within the haplogroup than those which are published.

We can see this easily in R-L48. Using all 84 non-repeating markers in Table A1 for which we have mutation rates, the TMRCA is 0.213/(2.32 x10^−3^) = 85 generations. Using the 23 markers in R-L48 for which mutational activity is reduced, one obtains only 32 generations to the coalescent. Using the 61 “healthy” markers one obtains 98 generations. The last estimate is the most likely to be correct – depending on what haplotypes were most strongly represented in the father-son sample from which the mutation times were calculated.

Another estimate of the coalescent can also be made by finding two men with the greatest genetic distance. We have two men in the L48 sample who only match on 65 out of 111 markers, and because many marker differences are 2-3 repeats, their “genetic (block) distance” is 63. The Clan Donald TMRCA calculator for pairs of men shows a fairly good peak at 118 generations for a 65 of 111 marker match. The coalescent time, taking into account the multistep differences, can be expected to be quite a bit earlier than this, and the 98 generations calculated from the variance quotient is a clear underestimate.

Dating E1b1b1c is also sensitive to the choice of markers in the variance test. Using all markers, it has a TMRCA of 269 generations. Using the “low variance” markers it is 206 generations and with the “healthy” ones it is 355 generations.

### Microsatellite life cycle

The Y-chromosome is generally regarded as the most degraded part of the human genome, because so little of it consists of functional genes and so much is “junk DNA” – repetitive motifs, degenerate relics of ancient X-genes and other sequences without apparent function. Surprisingly however, the human Y-chromosome has little variation, much less than elsewhere in the genome (Wilson Sayres *et al.* 2014). However it does undergo occasional “unique event polymorphisms”– deletions and, insertions. In non-active regions these can delete or disrupt microsatellites and change large numbers of single bases. A 2kb deletion originally defined the J haplogroup. Deletions are implicated in the formation of some subclades of R1b, and, we suspect, within a number of other haplogroups.

Here we are more concerned with small-scale local events – the “slippage” mechanism (originally proposed in [11]), which causes regular microsatellite polymorphisms; and point mutations, which regularly change the performance of microsatellites. A consensus has developed, following [12], [13] and [14], that microsatellites have a “life cycle” in which they are born, expand to some stable value, decline and become inert. There is a fairly large literature on theoretical models of extension and decay of repeating microsatellite sequences ([15], [16])

These papers have postulated a combination of slippage to increase microsatellite lengths and point mutation to split sequences into smaller pieces. Several authors ([17], [3]) have used the term “microsatellite death” for loss of mutational activity, and suggest the build up of kurtosis in the distribution of allele values of a single marker is a sign that this position is approaching, though they do not describe death in any particular microsatellite. Kelkar *et al.* [18] investigated this microsatellite life cycle in primates, finding that microsatellites could last for tens or even hundreds of millions of years, concluding “substitutions were the cause for deaths of microsatellites of virtually all lengths”. Point mutations are stated to suppress slippage mutation rates by “disrupting local self-similarity”. But how does this work?

Now point mutations occur much less frequently than slippage, with a probability slightly more than 10^−8^ per generation per base [19]. As microsatellites have a total length typically less than 200 bases, point mutations in a typical microsatellite occur only once in a million generations or so. Because we can actually observe the decay of microsatellites in several stages during times of the order of 350 generations, point mutations cannot be responsible - though they are probably implicated. The degeneration must be a result of slippage or some other fast process.

<u>Slippage</u> occurs when the DNA strand “bunches up” in a small loop as it is spun out, rather as a cassette tape will bunch. The general description (eg [20]) is that the replicating strand and the template strand disassociate and then realign out of register, forming a loop in one of the strands. Slippages are common, especially short ones, but most of these are corrected by the various DNA correction systems [21]. The exact mechanism by which realignment occurs and how some slippages escape correction is not described. It would appear however that the polymerase proofreading system which follows the copying is fooled because the slippage must be an exact multiple of the motif length *m*, since it continues to encounter the correct DNA sequence after slippage, at least until the end of the microsatellite.

It is to be expected that slippages occur more frequently in longer repeat sequences, as there are more opportunities for rebinding or looping. There is a fairly large literature modelling this fact [22], [23], [24], [25]). Whittaker et al. [26, Figure 4] use a large database of father-son pairs to show that the mutation rate depends not just on the average length of different markers, but on different counts at the same locus. The mutation rate is low for counts of less than #10, and is apparently nonexistent for counts less than #7. This has importance for our inert markers.

### An example

Many of the microsatellites in use are “complex” – which usually means they are interrupted by point mutations To give an example of why a microsatellite might become inert, the following sequence (taken from [27]) shows the complex marker DYS552 in an individual from E haplogroup:

> . . . ..
>
> GGTGTTCTGATGAGGATAATT/TATAC/TATAC/
>
> TGTAC/TGTAC/TATAC/TATAC/TATAC/TATAC/
>
> TATAC/TATAC/TATAC/TATAC/TATAC/TATAC/
>
> TATAC/TATAC/TATAC/TATAC/CATAC/TATAC/
>
> CATAC/TATAC/TATAC/TATAC/CATAC/CATAC/
>
> TATAC/TATAC/TATAC/CATAC/TATAC/TATAC/
>
> TATAC/AACCAATTAATTAGCTGAGTATAATAA
>
> . . . . .

Within the beginning and end blocks are 33 pentabase motifs – one main motif TATAC with 26 repeats, and two other scattered motifs TGTAC, CATAC altered from the main by a single base, presumably due to two point mutations. Now under the International Society for Forensic Genetics (ISFG) and the guidelines of the US National Institute of Standards and Technology (NIST) the full count of 33 is taken to be the marker value, with all the different repeats included. This explains how the modal value of the marker count is maintained even as point mutations accumulate. Slippage mutations normally occur in the single continuous twelve base sequence [29], which is the only one long enough for conventional slippage to occur.

However it is clear in this example that at some time the irregular motifs have also been copied and lie within a few motifs of the original occurrence. This might have been due to (rare) 10 or 15 base slippages. In [28] it is show that short indels are normally very rare, slower than point mutations, but they occur with much greater frequency in repeat area “hotspots” of the genome, even when the repeats are not as regular as in tandem sequences.

When the corrupted motifs are duplicated similarly as some of the examples given in [30] they can further interrupt any remaining long “clean” sequences, causing them to be too short for rapid slippage changes to occur. Therefore the marker appears to be dead.

Does a microsatellite actually “die” or does it just become essentially inert, but capable of accidental repair? Whether or not a microsatellite can resume activity would depend on exactly what is preventing it from mutating by slippage. It is possible that a fortuitous deletion of a “dirty” section could restore marker polymorphism again.

Towards the end of their lives, perhaps “slow” microsatellites are moving in and out of the inert state due to fortuitous rare slippage indels. We have seen how DYS472 seems to have sprung “back to life” in E1b1b. Perhaps the term “microsatellite death” should be reserved for the outright annihilation of a microsatellite due to larger-scale mutations, the null result, rather than for loss of its polymorphism capability.

### Discussion and Conclusions

A number of authors have explored the interaction of slipping and point mutations in microsatellites, speculating on a “microsatellite life cycle” of birth, expansion, degeneration and death, , but as far as we know examples of the latter have not previously been demonstrated. Our larger-scale cross-sectional data shows that not only are mutation rates of microsatellites in different haplotypes very different, many have been undergoing a senescence process and a few are actually inert in some subclades. It is surprising that this has not been previously observed, but this may be because the large samples of over a hundred individuals from a single subclade necessary to demonstrate this have only recently become available. There is ample evidence that smaller local genetic events are affecting spreads and modal values of markers. As with other errors in replication, generally the effect is deleterious, slowing down the genetic activity of the microsatellites and sometimes leading to their “death”.

We have suggested that point mutations might be spread through the microsatellite by slippage insertions, which might explain the relatively rapid senescence and eventual death of markers within moderate numbers of generations. Sequencing of microsatellites on a selection of individuals, particularly those in populations with inert or degraded markers, should reveal what type of mechanism is actually involved.

An important outstanding research question is whether the different mutation rates of microsatellites can be deduced purely from their internal structure and process– as through some process such as the one we have just described – or whether more general mechanisms are regulating the mutation rate in different parts of the chromosome. This can be fairly easily resolved by establishing a library of microsatellite sequences, particularly those which have slowed or become inert. From this it may be possible to deduce exactly what mechanisms determine the “speed” of microsatellite mutation – to be added to the usual microsatellite mutation rate determinants of motif length and number of repeats.

While the phenomenon of marker senescence undermines one of the traditional uses of microsatellites in dating haplogroups, this is not fatal as plenty of markers maintain their viability within each haplotype, it is simply a case of excluding the degraded ones from the analysis. The more complex picture presented here in which different slippage rates are associated with modified microsatellite structures offers new opportunities for phenotyping through better understanding the complexities of microsatellite mutation.

## Acknowledgements

I thank Professor Mark Jobling, University of Leicester, for commenting on a draft of this paper, and in particular for bringing up the issue of short allele counts in determining slow mutation rates.

## Websites

(all accessed May 2014)

Little, L (2007).

Distribution of allele frequencies

http://freepages.genealogy.rootsweb.ancestry.com/∼geneticgenealogy/yfreq.htm

Gene diversity (variance)

http://freepages.genealogy.rootsweb.ancestry.com/∼geneticgenealogy/67GD.htm

Family Tree DNA projects

https://www.familytreedna.com/public/r1b/

https://www.familytreedna.com/public/R1b1b1/

https://www.familytreedna.com/public/U106/

https://www.familytreedna.com/public/Chaplogroup/

https://www.familytreedna.com/public/R1aY-Haplogroup

https://www.familytreedna.com/public/E1b1a

Clan Donald TMRCA calculator

http://dna-project.clan-donald-usa.org/tmrca.htm

## SUPPLEMENTARY MATERIAL

The comparison for each variance *v_ij_* for marker *i* in sample *j* is with the estimated value *v_ij_* = p_i_/pav.v_j_*, where *v_j_* is the average variance for the sample (last row of table) and p_av_ is the average transition probability (last row, second column). If the variance is less than 10 per cent of the estimate it is taken as extremely low, less than 40 per cent low, more than 160 per cent high.

This comparison depends on having accurate transition probabilities. Changing the probabilities will alter the balance between “low” and “high” variances; however the variances marked as “extremely low” are mostly unaffected. Changing this high-low balance does not alter the fact that the variances are significantly different in different subclades, it just changes what is “normal” for the marker, like turning a tone knob on a radio.

We have used probabilities from Ballantyne *et al.* (2010) because their estimates are quite well correlated with our variances – which is probably because their father son pairs come from a similar, predominantly European, population. Burgarella and Navascues (2010) used wide-ranging populations and their fairly poor correlation with our sample probably reflects the different haplotypes they have used. However, they have estimates on about nine markers that match our sample variances much more closely than Ballantyne *et al*.

Another 20 markers appear to have different probabilities than either paper proposes. Any row that is mostly italics has the probability set too low, and if mostly bold – too high. This is not surprising, as both authors show very large confidence limits on their estimates.

We have not specifically investigated the role of mean marker allele length in determining the inter-haplotype differences – but there are many cases in which measured marker length is not the issue, with longer markers having a low variance.

**Table A1.**
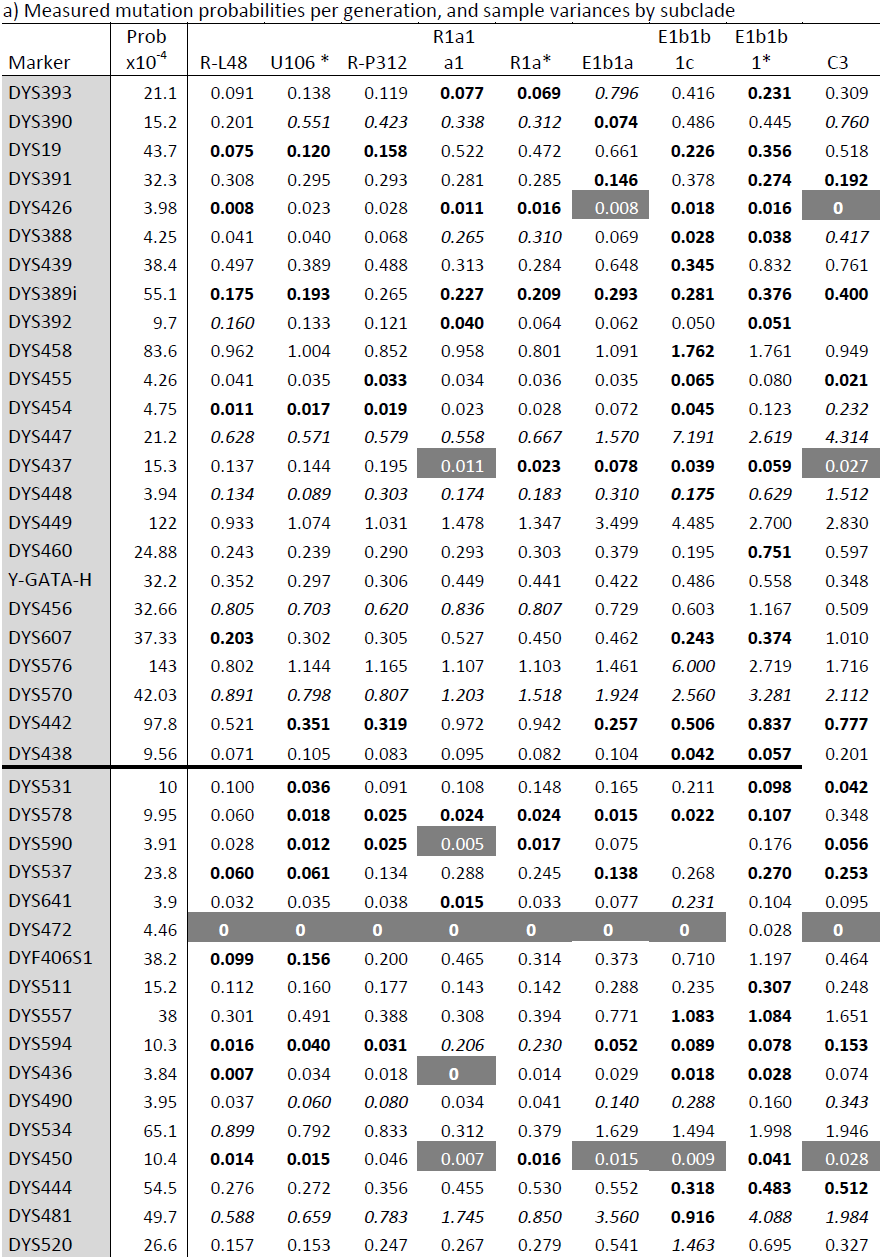
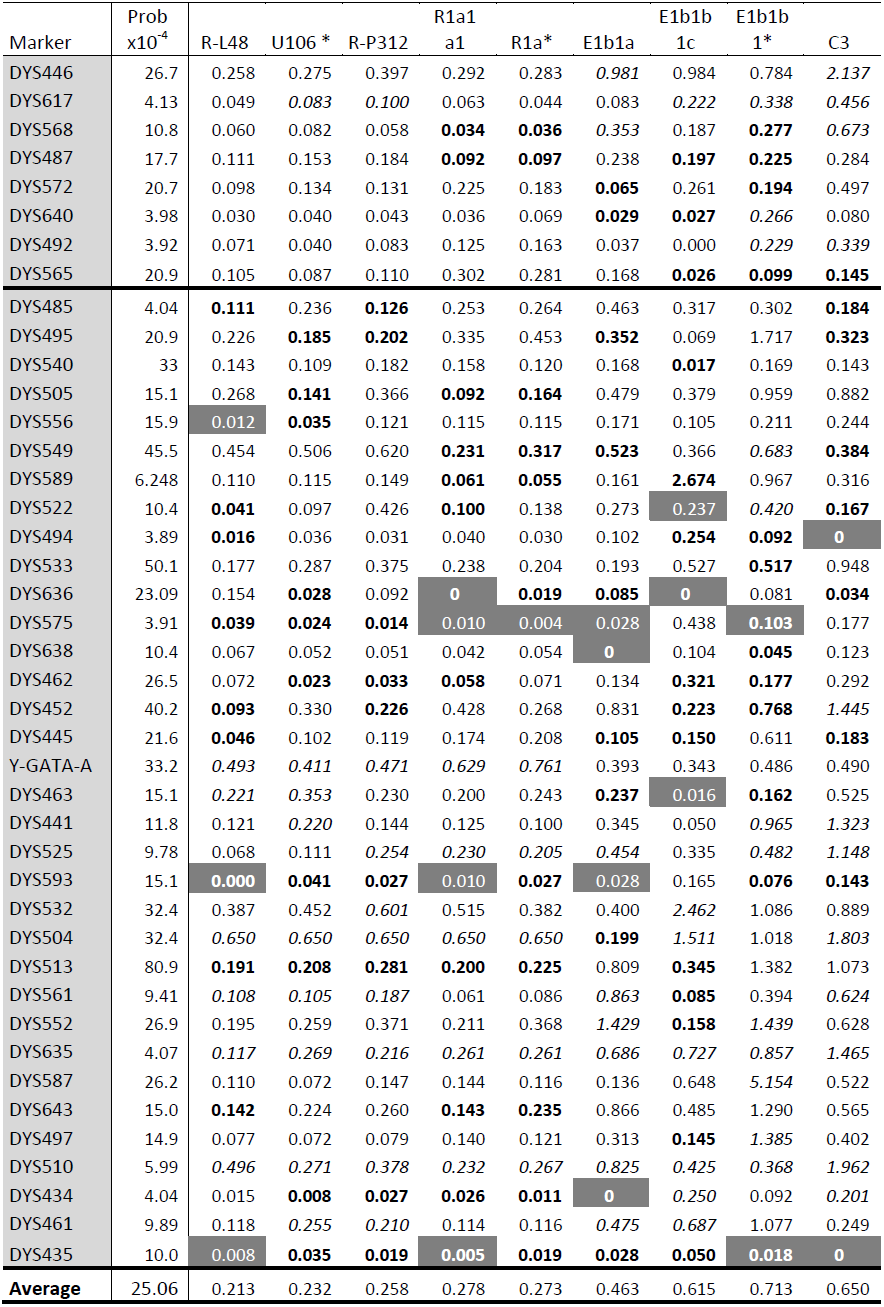
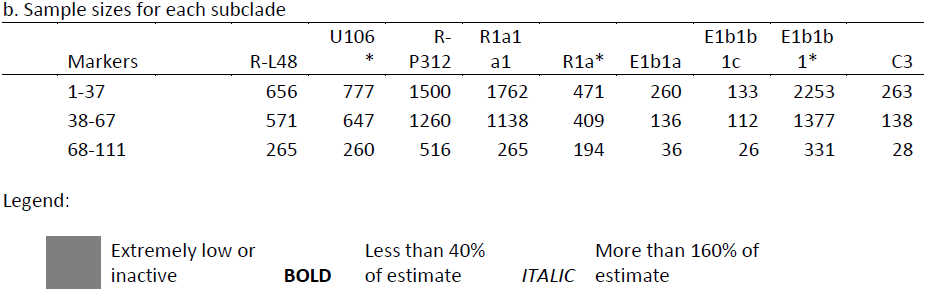
Marker probabilities and sample variances by selected subclades.

a) Measured mutation probabilities per generation, and sample variances by subclade
| Marker | Prob<br>x10<sup>-4</sup> |  |  |  | R1a1 |  | E1b1b |  | E1b1b |  |
| --- | --- | --- | --- | --- | --- | --- | --- | --- | --- | --- |
|  |  | R-L48 | U106 * | R-P312 | a1 | R1a* | E1b1a | 1c | 1* | C3 |
| DYS393 | 21.1 | 0.091 | 0.138 | 0.119 | **0.077** | **0.069** | 0.796 | 0.416 | **0.231** | 0.309 |
| DYS390 | 15.2 | 0.201 | 0.551 | 0.423 | 0.338 | 0.312 | **0.074** | 0.486 | 0.445 | 0.760 |
| DYS19 | 43.7 | **0.075** | **0.120** | **0.158** | 0.522 | 0.472 | 0.661 | **0.226** | **0.356** | 0.518 |
| DYS391 | 32.3 | 0.308 | 0.295 | 0.293 | 0.281 | 0.285 | **0.146** | 0.378 | **0.274** | **0.192** |
| DYS426 | 3.98 | **0.008** | 0.023 | 0.028 | **0.011** | **0.016** | **0.008** | **0.018** | **0.016** | **0** |
| DYS388 | 4.25 | 0.041 | 0.040 | 0.068 | 0.265 | 0.310 | 0.069 | **0.028** | **0.038** | 0.417 |
| DYS439 | 38.4 | 0.497 | 0.389 | 0.488 | 0.313 | 0.284 | 0.648 | **0.345** | 0.832 | 0.761 |
| DYS389i | 55.1 | **0.175** | **0.193** | 0.265 | **0.227** | **0.209** | **0.293** | **0.281** | **0.376** | **0.400** |
| DYS392 | 9.7 | 0.160 | 0.133 | 0.121 | **0.040** | 0.064 | 0.062 | 0.050 | **0.051** |  |
| DYS458 | 83.6 | 0.962 | 1.004 | 0.852 | 0.958 | 0.801 | 1.091 | **1.762** | 1.761 | 0.949 |
| DYS455 | 4.26 | 0.041 | 0.035 | **0.033** | 0.034 | 0.036 | 0.035 | **0.065** | 0.080 | **0.021** |
| DYS454 | 4.75 | **0.011** | **0.017** | **0.019** | 0.023 | 0.028 | 0.072 | **0.045** | 0.123 | 0.232 |
| DYS447 | 21.2 | 0.628 | 0.571 | 0.579 | 0.558 | 0.667 | 1.570 | 7.191 | 2.619 | 4.314 |
| DYS437 | 15.3 | 0.137 | 0.144 | 0.195 | 0.011 | **0.023** | **0.078** | **0.039** | **0.059** | 0.027 |
| DYS448 | 3.94 | 0.134 | 0.089 | 0.303 | 0.174 | 0.183 | 0.310 | **0.175** | 0.629 | 1.512 |
| DYS449 | 122 | 0.933 | 1.074 | 1.031 | 1.478 | 1.347 | 3.499 | 4.485 | 2.700 | 2.830 |
| DYS460 | 24.88 | 0.243 | 0.239 | 0.290 | 0.293 | 0.303 | 0.379 | 0.195 | **0.751** | 0.597 |
| Y-GATA-H | 32.2 | 0.352 | 0.297 | 0.306 | 0.449 | 0.441 | 0.422 | 0.486 | 0.558 | 0.348 |
| DYS456 | 32.66 | 0.805 | 0.703 | 0.620 | 0.836 | 0.807 | 0.729 | 0.603 | 1.167 | 0.509 |
| DYS607 | 37.33 | **0.203** | 0.302 | 0.305 | 0.527 | 0.450 | 0.462 | **0.243** | **0.374** | 1.010 |
| DYS576 | 143 | 0.802 | 1.144 | 1.165 | 1.107 | 1.103 | 1.461 | 6.000 | 2.719 | 1.716 |
| DYS570 | 42.03 | 0.891 | 0.798 | 0.807 | 1.203 | 1.518 | 1.924 | 2.560 | 3.281 | 2.112 |
| DYS442 | 97.8 | 0.521 | **0.351** | **0.319** | 0.972 | 0.942 | **0.257** | **0.506** | **0.837** | **0.777** |
| DYS438 | 9.56 | 0.071 | 0.105 | 0.083 | 0.095 | 0.082 | 0.104 | **0.042** | **0.057** | 0.201 |
| DYS531 | 10 | 0.100 | **0.036** | 0.091 | 0.108 | 0.148 | 0.165 | 0.211 | **0.098** | **0.042** |
| DYS578 | 9.95 | 0.060 | **0.018** | **0.025** | **0.024** | **0.024** | **0.015** | **0.022** | **0.107** | 0.348 |
| DYS590 | 3.91 | 0.028 | **0.012** | **0.025** | 0.005 | **0.017** | 0.075 |  | 0.176 | **0.056** |
| DYS537 | 23.8 | **0.060** | **0.061** | 0.134 | 0.288 | 0.245 | **0.138** | 0.268 | **0.270** | **0.253** |
| DYS641 | 3.9 | 0.032 | 0.035 | 0.038 | **0.015** | 0.033 | 0.077 | 0.231 | 0.104 | 0.095 |
| DYS472 | 4.46 | **0** | **0** | **0** | **0** | **0** | **0** | **0** | 0.028 | **0** |
| DYF406S1 | 38.2 | **0.099** | **0.156** | 0.200 | 0.465 | 0.314 | 0.373 | 0.710 | 1.197 | 0.464 |
| DYS511 | 15.2 | 0.112 | 0.160 | 0.177 | 0.143 | 0.142 | 0.288 | 0.235 | **0.307** | 0.248 |
| DYS557 | 38 | 0.301 | 0.491 | 0.388 | 0.308 | 0.394 | 0.771 | **1.083** | **1.084** | 1.651 |
| DYS594 | 10.3 | **0.016** | **0.040** | **0.031** | 0.206 | 0.230 | **0.052** | **0.089** | **0.078** | **0.153** |
| DYS436 | 3.84 | **0.007** | 0.034 | 0.018 | **0** | 0.014 | 0.029 | **0.018** | **0.028** | 0.074 |
| DYS490 | 3.95 | 0.037 | 0.060 | 0.080 | 0.034 | 0.041 | 0.140 | 0.288 | 0.160 | 0.343 |
| DYS534 | 65.1 | 0.899 | 0.792 | 0.833 | 0.312 | 0.379 | 1.629 | 1.494 | 1.998 | 1.946 |
| DYS450 | 10.4 | **0.014** | **0.015** | 0.046 | 0.007 | **0.016** | **0.015** | **0.009** | **0.041** | 0.028 |
| DYS444 | 54.5 | 0.276 | 0.272 | 0.356 | 0.455 | 0.530 | 0.552 | **0.318** | **0.483** | **0.512** |
| DYS481 | 49.7 | 0.588 | 0.659 | 0.783 | 1.745 | 0.850 | 3.560 | **0.916** | 4.088 | 1.984 |
| DYS520 | 26.6 | 0.157 | 0.153 | 0.247 | 0.267 | 0.279 | 0.541 | 1.463 | 0.695 | 0.327 |

| Marker | Prob<br>x10<sup>-4</sup> | R-L48 | U106 * | R-P312 | R1a1<br>a1 | R1a* | E1b1a | E1b1b<br>1c | E1b1b<br>1* | C3 |
| --- | --- | --- | --- | --- | --- | --- | --- | --- | --- | --- |
| DYS446 | 26.7 | 0.258 | 0.275 | 0.397 | 0.292 | 0.283 | 0.981 | 0.984 | 0.784 | 2.137 |
| DYS617 | 4.13 | 0.049 | 0.083 | 0.100 | 0.063 | 0.044 | 0.083 | 0.222 | 0.338 | 0.456 |
| DYS568 | 10.8 | 0.060 | 0.082 | 0.058 | **0.034** | **0.036** | 0.353 | 0.187 | **0.277** | 0.673 |
| DYS487 | 17.7 | 0.111 | 0.153 | 0.184 | **0.092** | **0.097** | 0.238 | **0.197** | **0.225** | 0.284 |
| DYS572 | 20.7 | 0.098 | 0.134 | 0.131 | 0.225 | 0.183 | **0.065** | 0.261 | **0.194** | 0.497 |
| DYS640 | 3.98 | 0.030 | 0.040 | 0.043 | 0.036 | 0.069 | **0.029** | **0.027** | 0.266 | 0.080 |
| DYS492 | 3.92 | 0.071 | 0.040 | 0.083 | 0.125 | 0.163 | 0.037 | 0.000 | 0.229 | 0.339 |
| DYS565 | 20.9 | 0.105 | 0.087 | 0.110 | 0.302 | 0.281 | 0.168 | **0.026** | **0.099** | **0.145** |
| DYS485 | 4.04 | **0.111** | 0.236 | **0.126** | 0.253 | 0.264 | 0.463 | 0.317 | 0.302 | **0.184** |
| DYS495 | 20.9 | 0.226 | **0.185** | **0.202** | 0.335 | 0.453 | **0.352** | 0.069 | 1.717 | **0.323** |
| DYS540 | 33 | 0.143 | 0.109 | 0.182 | 0.158 | 0.120 | 0.168 | **0.017** | 0.169 | 0.143 |
| DYS505 | 15.1 | 0.268 | **0.141** | 0.366 | **0.092** | **0.164** | 0.479 | 0.379 | 0.959 | 0.882 |
| DYS556 | 15.9 | **0.012** | **0.035** | 0.121 | 0.115 | 0.115 | 0.171 | 0.105 | 0.211 | 0.244 |
| DYS549 | 45.5 | 0.454 | 0.506 | 0.620 | **0.231** | **0.317** | **0.523** | 0.366 | 0.683 | **0.384** |
| DYS589 | 6.248 | 0.110 | 0.115 | 0.149 | **0.061** | **0.055** | 0.161 | **2.674** | 0.967 | 0.316 |
| DYS522 | 10.4 | **0.041** | 0.097 | 0.426 | **0.100** | 0.138 | 0.273 | **0.237** | 0.420 | **0.167** |
| DYS494 | 3.89 | **0.016** | 0.036 | 0.031 | 0.040 | 0.030 | 0.102 | **0.254** | **0.092** | **0** |
| DYS533 | 50.1 | 0.177 | 0.287 | 0.375 | 0.238 | 0.204 | 0.193 | 0.527 | **0.517** | 0.948 |
| DYS636 | 23.09 | 0.154 | **0.028** | 0.092 | **0** | **0.019** | **0.085** | **0** | 0.081 | **0.034** |
| DYS575 | 3.91 | **0.039** | **0.024** | **0.014** | 0.010 | 0.004 | 0.028 | 0.438 | **0.103** | 0.177 |
| DYS638 | 10.4 | 0.067 | 0.052 | 0.051 | 0.042 | 0.054 | **0** | 0.104 | **0.045** | 0.123 |
| DYS462 | 26.5 | 0.072 | **0.023** | **0.033** | **0.058** | 0.071 | 0.134 | **0.321** | **0.177** | 0.292 |
| DYS452 | 40.2 | **0.093** | 0.330 | **0.226** | 0.428 | 0.268 | 0.831 | **0.223** | **0.768** | 1.445 |
| DYS445 | 21.6 | **0.046** | 0.102 | 0.119 | 0.174 | 0.208 | **0.105** | **0.150** | 0.611 | **0.183** |
| Y-GATA-A | 33.2 | 0.493 | 0.411 | 0.471 | 0.629 | 0.761 | 0.393 | 0.343 | 0.486 | 0.490 |
| DYS463 | 15.1 | 0.221 | 0.353 | 0.230 | 0.200 | 0.243 | **0.237** | **0.016** | **0.162** | 0.525 |
| DYS441 | 11.8 | 0.121 | 0.220 | 0.144 | 0.125 | 0.100 | 0.345 | 0.050 | 0.965 | 1.323 |
| DYS525 | 9.78 | 0.068 | 0.111 | 0.254 | 0.230 | 0.205 | 0.454 | 0.335 | 0.482 | 1.148 |
| DYS593 | 15.1 | **0.000** | **0.041** | **0.027** | 0.010 | **0.027** | **0.028** | 0.165 | **0.076** | **0.143** |
| DYS532 | 32.4 | 0.387 | 0.452 | 0.601 | 0.515 | 0.382 | 0.400 | 2.462 | 1.086 | 0.889 |
| DYS504 | 32.4 | 0.650 | 0.650 | 0.650 | 0.650 | 0.650 | **0.199** | 1.511 | 1.018 | 1.803 |
| DYS513 | 80.9 | **0.191** | **0.208** | **0.281** | **0.200** | **0.225** | 0.809 | **0.345** | 1.382 | 1.073 |
| DYS561 | 9.41 | 0.108 | 0.105 | 0.187 | 0.061 | 0.086 | 0.863 | **0.085** | 0.394 | 0.624 |
| DYS552 | 26.9 | 0.195 | 0.259 | 0.371 | 0.211 | 0.368 | 1.429 | **0.158** | 1.439 | 0.628 |
| DYS635 | 4.07 | 0.117 | 0.269 | 0.216 | 0.261 | 0.261 | 0.686 | 0.727 | 0.857 | 1.465 |
| DYS587 | 26.2 | 0.110 | 0.072 | 0.147 | 0.144 | 0.116 | 0.136 | 0.648 | 5.154 | 0.522 |
| DYS643 | 15.0 | **0.142** | 0.224 | 0.260 | **0.143** | **0.235** | 0.866 | 0.485 | 1.290 | 0.565 |
| DYS497 | 14.9 | 0.077 | 0.072 | 0.079 | 0.140 | 0.121 | 0.313 | **0.145** | 1.385 | 0.402 |
| DYS510 | 5.99 | 0.496 | 0.271 | 0.378 | 0.232 | 0.267 | 0.825 | 0.425 | 0.368 | 1.962 |
| DYS434 | 4.04 | 0.015 | **0.008** | **0.027** | **0.026** | **0.011** | **0** | 0.250 | 0.092 | 0.201 |
| DYS461 | 9.89 | 0.118 | 0.255 | 0.210 | 0.114 | 0.116 | 0.475 | 0.687 | 1.077 | 0.249 |
| DYS435 | 10.0 | **0.008** | **0.035** | **0.019** | **0.005** | **0.019** | **0.028** | **0.050** | **0.018** | **0** |
| Average | 25.06 | 0.213 | 0.232 | 0.258 | 0.278 | 0.273 | 0.463 | 0.615 | 0.713 | 0.650 |

b. Sample sizes for each subclade
| Markers | R-L48 | U106<br>* | R-P312 | R1a1<br>a1 | R1a* | E1b1a | E1b1b<br>1c | E1b1b<br>1* | C3 |
| --- | --- | --- | --- | --- | --- | --- | --- | --- | --- |
| 1-37 | 656 | 777 | 1500 | 1762 | 471 | 260 | 133 | 2253 | 263 |
| 38-67 | 571 | 647 | 1260 | 1138 | 409 | 136 | 112 | 1377 | 138 |
| 68-111 | 265 | 260 | 516 | 265 | 194 | 36 | 26 | 331 | 28 |

Legend:
|  |  |  |  |  |  |
| --- | --- | --- | --- | --- | --- |
|  | Extremely low or inactive | **BOLD** | Less than 40% of estimate | *ITALIC* | More than 160% of estimate |

